# An exact transformation for CNN kernel enables accurate sequence motif identification and leads to a potentially full probabilistic interpretation of CNN

**DOI:** 10.1101/163220

**Authors:** Yang Ding, Jing-Yi Li, Meng Wang, Xinming Tu, Ge Gao

## Abstract

**Motivation:** Convolutional neural network (CNN) has been widely used in functional motifs identification for large-scale DNA/RNA sequences. Currently, however, the only way to interpret such a convolutional kernel is a heuristic construction of a position weight matrix (PWM) from fragments scored highly by that kernel.

**Results:** Instead of using heuristics, we developed a novel, exact kernel-to-PWM transformation whose equivalency is theoretically proven: the log-likelihood of the resulting PWM generating any DNA/RNA sequence is exactly the sum of a constant and the convolution of the original kernel on the same sequence. Importantly, we further proved that the resulting PWM’s performance on sequence classification/regression can be exactly the same as the original kernel’s under popular CNN frameworks. In simulation, the exact transformation rivals or outperforms the heuristic PWMs in terms of classifying sequences with sequence- or structure-motifs. The exact transformation also faithfully reproduces the output of CNN models on real-world cases, while the heuristic one fails, especially on the case with little prior knowledge on the form of underlying true motifs. Of note, the time complexity of the novel exact transformation is independent on the number of input sequences, enabling it to scale well for massive training sequences.

**Availability:** Python scripts for the transformation from kernel to PWM, the inverted transformation from PWM to kernel, and a proof-of-concept for the maximum likelihood estimation of optimal PWM are available through https://github.com/gao-lab/kernel-to-PWM.

## Introduction

Effective *ab inito* mining sequence motifs from large number of sequences is critical to mechanistic delineation of how the sequences exert their functions (Mathelier *et al.*, 2016; Ray *et al.*, 2013; Nawrocki *et al.*, 2015). While classical methods are either computationally expensive and thus limited to low number of sequences only (e.g., MEME (Bailey *et al.*, 2009)), or with strong assumptions on input sequences (e.g. MEME-ChIP (Machanick and Bailey, 2011) assumes that all input sequences must be centered and have equal length, which might greatly bias the result when it handles sequences other than those coming from ChIP-Seq), researchers have discovered recently that convolutional neural networks (CNNs) are exceptionally good at handling such problem fast and accurately (Alipanahi *et al.*, 2015; Angermueller *et al.*, 2017; Quang and Xie, 2016; Zhou and Troyanskaya, 2015). Basically, a CNN uses a series of convolutional kernels as motif detectors to identify potential sequence patterns. Each of these kernels is a matrix which is used to scan the input sequences, i.e., being “align”ed to each position. For each such “alignment”, a score is computed by convoluting the kernel over the aligned sequence fragment (LeCun *et al.*, 1989; Cotter *et al.*, 2011). Intuitively, the larger the score, the more “similar” the kernel is to the specific sequence fragment it aligns to. However, it remains challenging to extract/interpret the particular sequence patterns learned by these convolutional kernels.

Alipanahi *et al.* proposed a heuristic transformation to convert a given kernel into a Position Weight Matrix, or PWM (Alipanahi *et al.*, 2015), which has been widely used to represent experimentally determined DNA-(Mathelier *et al.*, 2016) or RNA-binding motifs (Ray *et al.*, 2013). Briefly, it transforms a kernel by (1) first stacking the kernel’s highly scoring sequence fragments from input sequences, and then (2) normalizing nucleotide counts per position to obtain the final PWM. Although the heuristic transformation has been widely used (e.g., Angermueller *et al.*, 2017; Kelley *et al.*, 2016; Ben-Bassat *et al.*, 2018), it is not guaranteed in theory (see Discussion for more details) and even fails in certain practical cases (see the AddGene case in Section Results later) to classify sequences as accurately as the kernels do.

In this paper, we proposed a novel, fast, exact kernel-to-PWM transformation (Figure 1(a)) to address this problem. We demonstrated, both theoretically and empirically, that the transformed PWM is capable of classifying/regressing sequences in exactly the same way as the original kernel (Figure 1(b)), indicating that this kernel interpretation is the most accurate among all possible interpretations. Further comparison to the heuristic transformation in both simulations and real-world cases showed that, when used to calculate PWM-based log-likelihood as a surrogate for convolutional output on the same input sequence (Figure 1(b)), the exact PWMs rival or greatly outperform the heuristic PWMs at detecting both sequence- and structure-motifs. Of note, our transformation also provides a direct and accurate probabilistic interpretation for the kernels used for motif detection (Zuallaert *et al.*, 2018). A Python implementation of the transformation is freely available online at https://github.com/gao-lab/kernel-to-PWM.

**Fig. 1.**
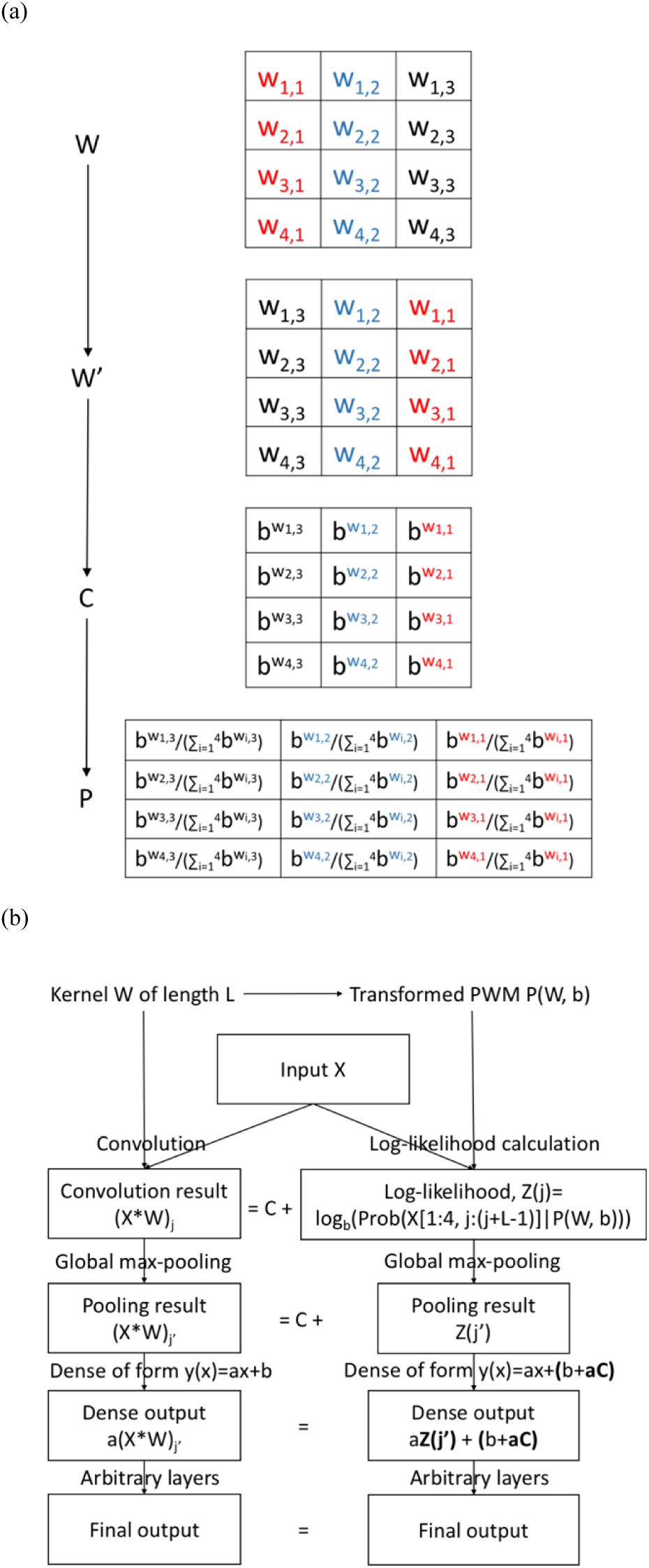
The transformation from kernel to PWM. (a) Steps of the trans-formation. Elements were colored to signify the flipping step (W -> W’). (b) The transformed PWM is capable of classifying/regressing sequences in exactly the same way as the original kernel, because it can (with parameters of succeeding layers adjusted accordingly) produce the same output as that the kernel produces. For simplicity, here we show this on a popular CNN structure with a single kernel only, but it can be easily extended to other popular structures (see the proof of Theorem 2 in Supplementary Notes) and the case with multiple kernels.

### The exact kernel-to-PWM transformation

Below we describe the transformation itself, and its interpretation. Detailed proofs of theorems and corollaries can be found in the Supplementary Notes.

#### The transformation

The five-step transformation, as illustrated in Figure 1(a), is described below (with all coordinates one-based; also see Algorithm 1):

1. Assume that the kernel to be transformed is a 4-by-L matrix W, where L is the length of this kernel. The element of W at the ith row and jth column is denoted as w_i,j_.
2. Choose an arbitrary base of logarithm, b (b > 1), for the log-likelihood calculation. There’s no further restriction on the choice of b.
3. Flip W along the second (position) axis to obtain the flipped kernel W’: w’_i,j_ = w_i,L-j+1_ for all i from {1, 2, 3, 4} and all j from {1, 2, …, L}.
4. Replace each w’_i,j_ with 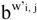 to obtain the exponentially transformed kernel C; in other words, 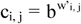 for all I from {1, 2, 3, 4} and all j from {1, 2, …, L}.
5. Normalize C in a column-wise manner by dividing each column by its sum, resulting in the PWM P(W, b): (P(W, b))_i,j_ = c_i,j_ / (c_1,_ j + c_2,j_ + c_3,j_ + c_4,j_) for all i from {1, 2, 3, 4} and all j from {1, 2,…, L}.

##### Algorithm 1: the transformation from kernel to PWM

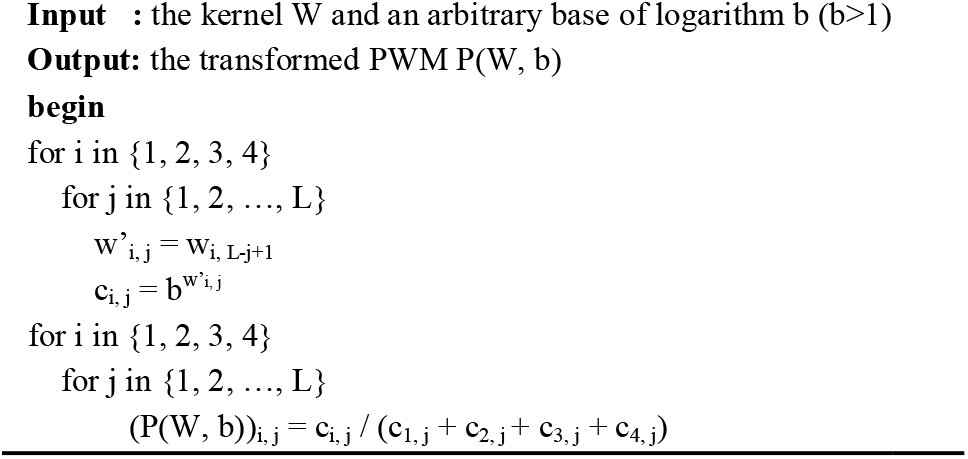

Then as stated by Theorem 1 (see Supplementary Notes for proof), for any b > 1 and any given sequence X no shorter than W, we have:

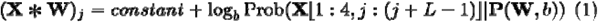

where * denotes the convolution operator that does not consider border-crossing cases (i.e., W must fall completely within X; see Supplementary Notes for details). In other words, the sequence’s convolution by W is exactly the sum of a constant and the log-likelihood of the PWM transformed on X. We also noted that the time complexity of the exact transformation (Algorithm 1) is O(L) and does not depend on the sequence datasets at all, making it very fast and easy to use.

Of note, the transformed PWM is thus capable of cl ssifying/regressing sequences in exactly the same way as the original kernel, because Equation (1) makes it possible to replace the output of convolutional layer with the corresponding log-likelihoods, while keeping the outputs of subsequent layers unchanged by a simple parameter tuning of immediately succeeding layers (see Figure 1(b) for a schematic explanation on a popular CNN structure; also see the row “Derivation of model parameters for our PWMs” in Supplementary Table 2 for a d tailed explanation involving multiple input sequences and kernels, and the proof of Theorems 1 and 2 in Supplementary Notes for an even more detailed mathematical treatment). This also immediately leads to two useful applications: kernel interpretation and PWM reuse.

#### Interpreting a kernel using the exact transformation

The exact transformation maps (and thus interprets) a kernel to (as) an infinite set of PWMs parameterized by b only, because Theorem 1 always holds regardless of the specific b chosen. These PWMs ranges from “uniform-like” (i.e., filled with values near 0.25; this happens when a b that is extremely close to 1 is used) to “fixed” (i.e., filled with values near 0 or 1; this happens when a very large b is used), yet, as demon-strated by Corollary 1 (See Supplementary Notes), any of them is capable of regressing/classifying the input sequences in exactly the same manner as the original kernel does under popular CNN frameworks (e.g. those in (Alipanahi *et al.*, 2015; Quang and Xie, 2016; Zhou and Troyanskaya, 2015)). Therefore, the user may choose a specific PWM of interest based on prior biological knowledge without biasing the kernel interpretation.

If no such prior knowledge is available, an alternative from the statistician’s view is to find the optimal b using maximum a posteriori estimation (MAP estimation). The exact form depends on the model structure and underlying assumptions, and should be treated on a case-by-ca e basis. As a proof-of-concept, we assumed a uniform distribution of b (thus making MAP estimation equivalent to maximum likelihood estimation), and then deduced and implemented in our repository the following estimate *q*(*b*|*X*^(1)^, …, *X*^(*n*)^,*W*^(1)^, … *W*^(*k*)^ a popular CNN framework in which an input sequence is fed to convolution, linear or ReLU activation, global max-pooling, linear function, and finally arbitrary functions (see Section 5 of Supplementary Notes for technical details):

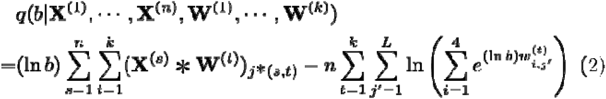

where X^(s)^ are the n indexed input sequences, W^(t)^ the k indexed kernels, L the length of each kernel, j*(s, t) the starting coordinate of the max-scored fragment by W^(t)^ on X^(s)^, and e the base of the natural logarithm.

#### Reusing a PWM in CNN models

Corollary 2 (see Supplementary Notes for details) guarantees that, from each PWM (with the restriction that no 0’s or 1’s are present), a kernel -- that is capable of classifying/regressing sequences in exactly the same manner as the PWM -- can be generated by the following steps (Figure 2 and Algorithm 2):

**Fig. 2.**
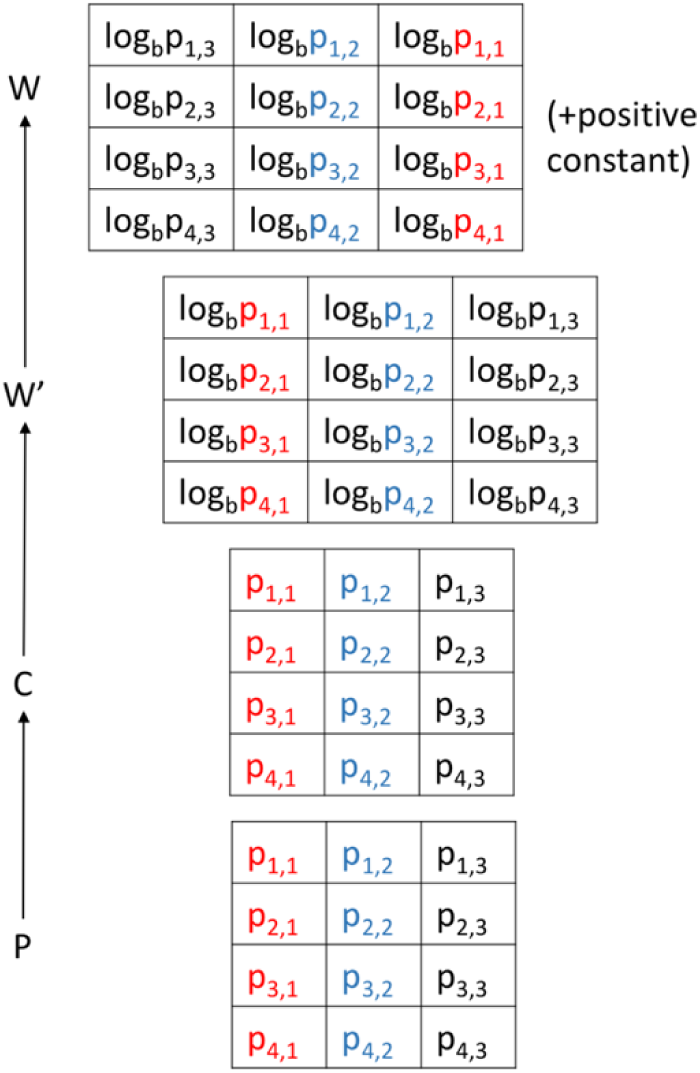
The (back) transformation from PWM to kernel. Elements were colored to signify the flipping step (W’->W). Note that the normalization step is skipped, and thus the C matrix is identical to P.

1. Similar to the transformation above, assume that the PWM to be transformed is a 4-by-L matrix P, where L is the length of this PWM. The element of P at the ith row and jth column is denoted as p_i,j_.
2. Choose an arbitrary base of logarithm, b (b>1);
3. Skip step 5 (normalization) by taking C = P;
4. Invert step 4 to obtain W’ (i.e., set w’_i,j_ = log_b_ c_i,j_);
5. Invert step 3 to obtain the transformed kernel W (i.e., set w_i,j_ = w’_i, L-j+1_);
6. If ReLU activation is to be used, add to the transformed kernel a positive shift sufficiently large to make all elements nonnegative.

##### Algorithm 2: the (back)-transformation from PWM to kernel

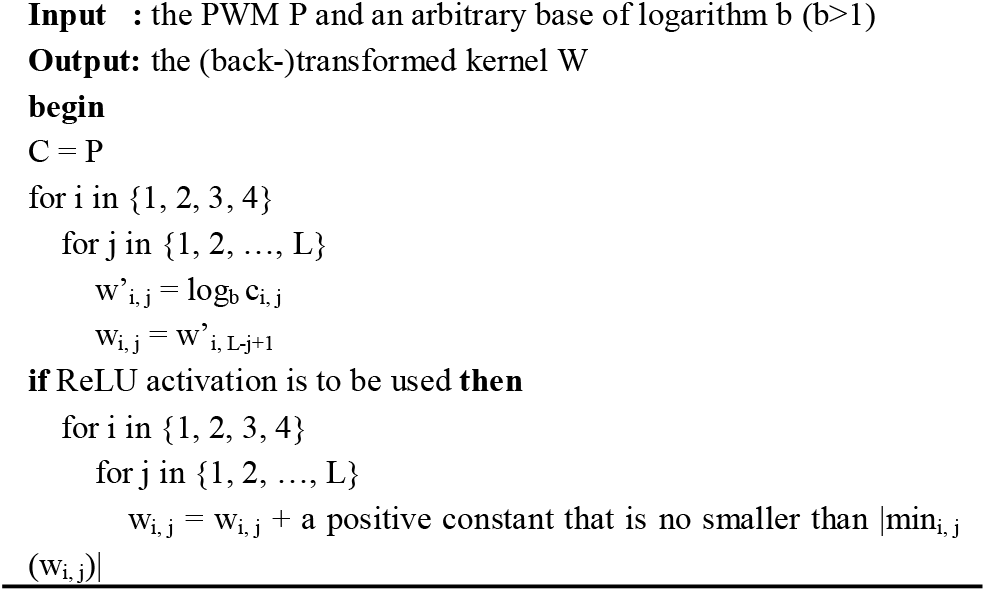

### Empirical results confirmed the outperformance of the exact transformation over the heuristic one at motif detection

Despite all the useful features above, it is still in doubt that the transformed PWMs can classify/regress sequences more accurately than PWMs transformed by other methods (specifically, the heuristic transformation proposed by Deepbind (Alipanahi *et al.*, 2015)).

In this section, we confirmed the outperformance of the exact transformation by both simulation and real-world cases. Briefly, for each trained CNN model in each case we obtained the two types of PWMs by both transformations, and examined which transformation’s PWM based log-likelihood can classify/regress the sequence (Figure 1(b)) better.

We note readers that, for PWMs from the exact transformation, parameters of layers succeeding the convolutional layer can be shifted/scaled accordingly to make the final output unchanged (see Figure 1(b) and proof of Theorem 2 in Supplementary Notes); in this way we do not need stochastic gradient descent to find the optimal parameters. We tuned parameters for these PWMs in this way for the two real-world cases; for the simulation case, stochastic gradient descent tends to work well enough already and we did not tune parameters in this way. For PWMs from the heuristic transformation, there’s no such known easy way of tuning parameters, and we need to use stochastic gradient descent to find the optimal parameters of these subsequent layers.

#### The exact transformation rivaled or outperformed the heuristic one at classifying simulated sequences with known sequence- and structure-motifs

In the simulation, we first extracted all motifs from known sequence (JASPAR (Mathelier *et al.*, 2016)) and structure (Rfam (Nawrocki *et al.*, 2015)) motif databases. For each motif, we compared the two transformations by the following pipeline (see Supplementary Table 1 for technical details):

1. We simulated positive and negative sequences, and trained a CNN model to classify these sequences;
2. We extracted the kernels and transformed each kernel into a PWM by either (1) the exact transformation or (2) the heuristic transformation from Deepbind (as in Section 10.2 “Sequences Logos” of its original Supplementary note);
3. We evaluated which transformation would better classify sequences (Figure 1(b)) by comparing their area under ROC curves (AUROC) and area under PR curves (AUPRC) across the entire dataset.

We then examined the overall outperformance by performing a Wil-coxon signed-rank test for both AUPRC and AUROC (NULL hypothesis: the mean AUC of the exact transformation is equal to or less than the mean AUC of the heuristic transformation). As expected, we found that the exact transformation significantly outperformed the heuristic one for structure-oriented Rfam motifs (Wilcoxon signed-rank test p-value < 2.2e-16 for both AUPRC and AUROC, and the mean absolute difference in AUC is 0.049 for AUPRC and 0.046 for AUROC), while almost rivaled for sequence-oriented JASPAR motifs (Wilcoxon signed-rank test p-value < 2.2e-16 for AUPRC and =1.053e-09 for AUROC, though the mean absolute difference in AUC is around 0.001).

#### The exact transformation can better recover the motif detection performance of real-world CNN models

We further demonstrated by two real-world cases that the exact transformation can better recover the motif detection performance of real-world CNN models.

The first case (Deepbind itself) trained a separate CNN-based motif detector for each of all the 461 real-world ChIP-Seq and SELEX motifs that are not deprecated by Deepbind’s paper (Alipanahi *et al.*, 2015). For each of these motifs, we compared the two transformations by the following pipeline (with the technical details available in Supplementary Table 2): (1) get the pre-trained CNN model from Deepbind (http://tools.genes.toronto.edu/deepbind/deepbind-v0.11-linux.tgz); (2) use this model to generate the benchmark dataset, with input sequences also from Deepbind (http://tools.genes.toronto.edu/deepbind/nbtcode/nbt3300-supplementary-software.zip); (3) transform the model’s kernels by both transformations; and (4) test for each transformation whether the modified model, where convolutional outputs are replaced by the corresponding log-likelihoods, replicates the CNN model’s performance after parameter tuning (for the exact transformation) / retraining (for the heuristic transformation).

We compared MAPE and MSE between the exact transformation and the heuristic one across all these motifs (Figure 3). Consistent with the previous subsection, the exact transformation outperformed Deepbind’s transformation on real datasets.

**Fig. 3.**
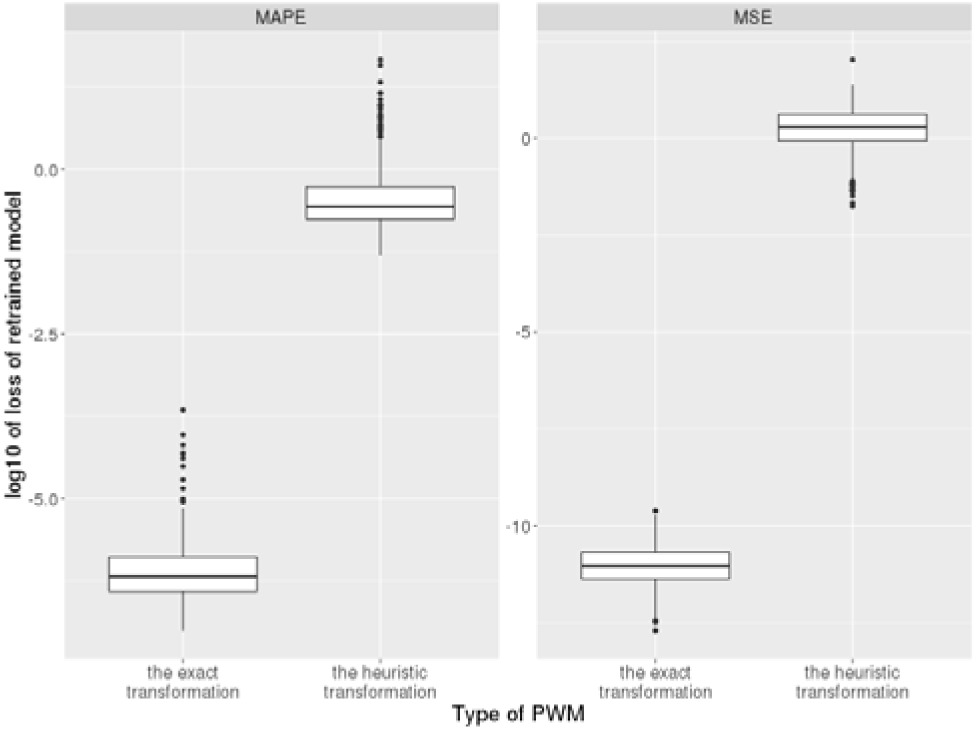
The exact transformation outperformed the heuristic one in exemplar motifs of Deepbind’s original work. For each of these motifs, the log-likelihoods from the two transformations were tested on how close they could approximate the output of original CNN model trained by Deepbind itself (by both mean absolute percentage error (MAPE) and mean squared error (MSE)). Codes for reproducing this plot are available on the Github repository.

The second case trained a CNN to predict the lab-of-origin of a plasmid by its sequence only (Nielsen and Voigt, 2018). Similar to the pipeline above, to compare the two transformations on this case, we (1) requested the dataset from AddGene, (2) obtained the PWMs by both transformations, and finally (3) computed the validation accuracy of classification for the two transformations (details are available through Supplementary Table 3).

The retrained CNN model has a validation accuracy of 47.49%, which appears to replicate the one reported by Nielsen and Voigt (48%); the small deviation might result from the augmentation of the requested dataset both in sample size (from 36764 to 64149) and number of different labels (from 827 to 1366) since publication. Based on this successfully retrained CNN model, our expectation still holds: PWMs from the exact transformation achieved almost the same validation accuracy (47.34%), while the heuristically transformed PWMs only have a validation accuracy of 1.61% even after training with stochastic gradient descent. This stark difference immediately invalidated the possibility that the heuristic transformation always represents the underlying true motif.

## Discussion

Convolutional neural network (CNN) have been widely used in learning regulatory sequence codes (Zhou and Troyanskaya, 2015; Kelley *et al.*, 2016), identifying binding sites (Alipanahi *et al.*, 2015; Quang and Xie, 2016), and calling various functional elements (Umarov and Solovyev, 2017) on large amount of DNA/RNA sequences. The CNN convolves, or “scans”, these sequences with multiple kernels. While these kernels have been widely thought to be sort of representation for underlying functional elements, only a heuristic transformation (Alipanahi *et al.*, 2015) was proposed for converting kernels to position weight matrices (PWM), a commonly used representation of sequence motifs.

Currently, convolutional kernels used for mining nucleotide sequences are just real matrices that need not conform to probabilistic restrictions of PWMs (e.g., each element should be a real number within [0, 1]). This discrepancy makes it heuristically difficult to accurately interpret kernels as PWMs. We have found for the first time that the kernel, together with a logarithm base, can be transformed into a PWM with a log-likelihood that is the sum of a constant and the kernel’s convolution. The mathematical solidity of this method connects intimately the computational field of CNNs and the biological field of functional elements, and both empirical simulations and testing on real-world cases have further demonstrated the superiority of the exact transformation to the heuristic one. Of note, the time complexity of the exact transformation is O(L) for a kernel of length L and does not depend on the number of input sequences, rendering it scaling well for models with thousands (or even millions) of training sequences; in contrast, the heuristic transformation sums fragments of all training sequences passing activation (Alipanahi *et al.*, 2015) and doesn’t scale well for large amount of data..

The failure of the heuristic transformation could be partially due to its inherent setup of optimization. From a perspective of maximum likeli-hood estimation (MLE), the resulting PWM maximizes the probability of observing highly scoring fragments given the PWM itself, but does not take into account the kernel itself; i.e., for the t-th kernel W^(t)^ it tries to find the PWM P^(t)^ that maximizes the following conditional probabili y (using notations from the MLE above), but without considering the specific value of the kernel to transform:

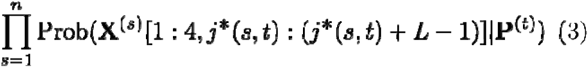

If we further impose the condition that the specific value of the kernel is also observed and should not be biased, we will end up with a combination of such MLE with the exact transformation; that is, it will try to find the PWM P^(t)^ that maximizes the following conditional probabili y and that is subject to the following restriction:

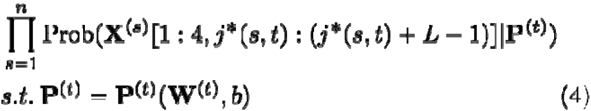

This could explain partially why the exact transformation outperformed Deepbind’s in some cases, especially when we’d like to recover the performance of a CNN model whose output is continuous.

Beyond PWMs, the development of this exact transformation compelled us to seek for similar transformations of more complex motif representations. In fact, PWMs are essentially 0-order Markov chains, and it is thus very natural to find the deep-learning counterpart of Markov chains with 1- or higher orders. Such model might have unexpected performance and efficiency on learning complex motifs from real biological datasets.

We’d like to highlight that our transformation also provides a new statistical view to understand the convolutional neural network. In this transformation, the score of the convolution on the given sequence is (up to a constant difference) equal to the log-probability of PWM to generate the same sequence. Based on this equivalence, both the subsequent pooling layer and the dense layer can find their exact, statistical interpretation: the pooling layer just computes (for global max-pooling) the largest or (for global average-pooling) the mean log-probability, and the dense layer computes a weighted joint log-probability ratio from different kernels (as the coefficient of kernels might have different signs). Our results well suggest a full probabilistic model for convolutional neural network, which enables a rational design and optimization for customized tasks in near future.

## Funding

This work was supported by funds from the National Key Research and Development Program (2016YFC0901603), the China 863 Program (2015AA020108), as well as the State Key Laboratory of Protein and Plant Gene Research and the Beijing Advanced Innovation Center for Genomics (ICG) at Peking University. The research of G.G. was supported in part by the National Program for Support of Top-notch Young Professionals. Part of the analysis was performed on the Computing Platform of the Center for Life Sciences of Peking University, and we thank Dr. Fangjin Chen for his help.

## Conflict of Interest

none declared.

