## Supplementary Notes for "An exact transformation for CNN kernel enables accurate sequence motif identification and leads to a potentially full probabilistic interpretation of CNN"

#### 1 Definitions

**Common 1.** *Throughout this text, matrices are denoted by boldface capital letters, e.g.,* $\mathbf{A} = (a_{i,j})$*, and vectors are denoted by boldface lowercase letters, e.g.,* $\mathbf{a} = (a_i)$*. All vectors are assumed to be column vectors. All subscripts are one-based.*

*In the proof we need to describe the submatrix of a given matrix. We use* $\mathbf{A}[i_1 : i_2, j_1 : j_2], i_1 \leqslant i_2, j_1 \leqslant j_2$ *to describe the submatrix of* $\mathbf{A}$ *generated by extracting rows* $i_1, i_1 + 1, \cdots, i_2$ *and columns* $j_1, j_1 + 1, \cdots, j_2$*. Sometimes, the submatrix columns are not consecutive in the given matrix; in this case, we use* $\mathbf{A}[[i_1, i_2, \cdots, i_{H_1}], [j_1, j_2, \cdots, j_{H_2}]]$ *to describe the submatrix of* $\mathbf{A}$ *generated by first extracting rows* $i_1, i_2, \cdots, i_{H_1}$ *and then extracting columns* $j_1, j_2, \cdots, j_{H_2}$ *(note that the orders of these row and column indices matter here). The enumeration* $[i_1, i_2, \cdots, i_H]$ *can also be replaced with* $order(i_1, i_2, \cdots, i_H)$*, i.e., an ordering of the set consisting of i's; in this case, the ordering is assumed to be ascending.*

*For a matrix* $\mathbf{A}$ *and a scalar* $b$*, we assume* $\mathbf{A} + b$ *is an element-wise addition, i.e.* $(\mathbf{A} + b)_{i,j} := a_{i,j} + b$*.*

*The vector* $\mathbf{1}_a$ *is a column vector of length a filled with 1.*

**Common 2.** *We use* $\mathbf{W}$ *as the matrix representation of an arbitrary kernel. It always has 4 rows, and the number of its columns is denoted by* $L (L > 0)$*.*

**Common 3.** *We use* $\mathbf{X}$ *as the matrix representation of an arbitrary input sequence. Similar to kernels, it always has 4 rows. The number of its columns is denoted by* $N$*. In the current problem setting, we only consider cases where input sequences are not shorter than kernels, and thus we assume that* $N \geqslant L$*.*

*We use one-hot encoding to represent an arbitrary input sequence using a matrix. Specifically, for any sequence* $(s_1, \cdots, s_N)$ *with* $s_j \in \{A, C, G, T\}, \forall j \in 1, \cdots, N$*, its matrix representation* $\mathbf{X}$ *could be written explicitly in the following form:*

$$
\begin{aligned}
x_{1j} &:= \begin{cases} 1 & s_j = A \\ 0 & otherwise \end{cases} \\
x_{2j} &:= \begin{cases} 1 & s_j = C \\ 0 & otherwise \end{cases} \\
x_{3j} &:= \begin{cases} 1 & s_j = G \\ 0 & otherwise \end{cases} \\
x_{4j} &:= \begin{cases} 1 & s_j = T \\ 0 & otherwise \end{cases}
\end{aligned} \tag{1}
$$

**Common 4.** *For the sake of convenience we assume that, if $a_1 > a_2$, then the summation $\sum_{i=a_1}^{a_2} (x_i) = 0$.*

**Definition 1.** *We define $f(j|\mathbf{X}) : \{1, \cdots, N\} \rightarrow \{1, 2, 3, 4\}$ to describe the nucleotide identity of the $j$-th nucleotide of the sequence that is represented by the matrix $\mathbf{X}$. Specifically, we have*

$$
f(j|\mathbf{X}) := i \ satisfying \ x_{i,j} = 1 \tag{2}
$$

*Since each column of $\mathbf{X}$ has one and only one element being 1, this function is well-defined.*

**Definition 2.** *We define $\mathbf{P}(\mathbf{W}, b) : \mathbb{R}^{4\times L} \rightarrow \mathbb{R}^{4\times L}$ as the matrix representation of the PWM of the sequence profile transformed from the kernel $\mathbf{W}$ with base for logarithm $b > 0$. Details of the transformation are described explicitly in the main text.*

**Definition 3.** *We define $\mathbf{X} \ast \mathbf{W}$ as the discrete convolution with matrix $\mathbf{W}$ on matrix $\mathbf{X}$. Specifically, we have:*

$$
(\mathbf{X} \ast \mathbf{W})_j := \sum_{i=1}^{4} \sum_{j'=1}^{L} x_{i,j+j'-1} w_{i,L-j'+1} \tag{3}
$$

**Definition 4.** *We define $\mathrm{Prob}(\mathbf{X}|\mathbf{P}(\mathbf{W}, b))$ as the probability that the sequence represented by matrix $\mathbf{X}$ is generated from the sequence profile whose PWM is represented by the matrix $\mathbf{P}(\mathbf{W}, b)$ defined above.*

**Definition 5.** *We define the elementwise exponential* $b^{\mathbf{A}}$ *as:*

$$\left(b^{\mathbf{A}}\right)_{i,j} := b^{(\mathbf{A}_{i,j})} \tag{4}$$

**Definition 6.** *We define* $\text{threshold}(\mathbf{x}, z)$ *as:*

$$(\text{threshold}(\mathbf{x}, z))_i := \begin{cases} x_i, & \text{if } x_i > z \\ 0, & \text{otherwise} \end{cases} \tag{5}$$

*Specifically, we define* $(\mathbf{x})_+ := \text{threshold}(\mathbf{x}, 0)$.

### 2 Theorem 1 and its proof

**Theorem 1.** $\forall \mathbf{W}, b > 0$*, transform them into the PWM* $\mathbf{P} = \mathbf{P}(\mathbf{W}, b)$ *according to the algorithm described in the main text of this paper. Then,* $\forall \mathbf{X}$*, we have:*

$$(\mathbf{X} \ast \mathbf{W})_j = \log_b \text{Prob}(\mathbf{X}[1:4, j:(j+L-1)] | \mathbf{P}(\mathbf{W}, b)) + d(\mathbf{W}, b) \tag{6}$$

*where* $d(\mathbf{W}, b) := \sum_{j'=1}^{L} \log_b \left( \sum_{i'=1}^{4} b^{w_{i',L-j'}} \right)$ *is a constant that does not depend on* $\mathbf{X}$*, but entirely on* $\mathbf{W}$ *and* $b$*.*

*Proof.* By definition of PWM, the probability that a sequence is generated from a PWM is the product of generating probability of "the nucleotide at each position in that sequence" at the same position in this PWM. Mathematically, this is:

$$\begin{aligned} &\text{Prob}(\mathbf{X}[1:4, j:(j+L-1)] | \mathbf{P}(\mathbf{W}, b)) \\ =& \prod_{j'=1}^{L} (\mathbf{P}(\mathbf{W}, b)[f(j' | \mathbf{X}[1:4, j:(j+L-1)]), j']) \end{aligned} \tag{7}$$

Taking logarithm with base $b$ of both sides gives the following result:

$$\begin{aligned} &\log_b \text{Prob}(\mathbf{X}[1:4, j:(j+L-1)] | \mathbf{P}(\mathbf{W}, b)) \\ =& \log_b \prod_{j'=1}^{L} \left( (\mathbf{P}(\mathbf{W}, b))_{f(j' | \mathbf{X}[1:4, j:(j+L-1)]), j'} \right) \\ =& \sum_{j'=1}^{L} \log_b \left( (\mathbf{P}(\mathbf{W}, b))_{f(j' | \mathbf{X}[1:4, j:(j+L-1)]), j'} \right) \\ =& \sum_{j'=1}^{L} \log_b \left( (\mathbf{P}(\mathbf{W}, b))_{f(j+j'-1 | \mathbf{X}), j'} \right) \end{aligned} \tag{8}$$

On the other hand, for the convolution we have:

$$
\begin{aligned}
&(\mathbf{X} * \mathbf{W})_j \\
=&\sum_{i=1}^{4} \sum_{j'=1}^{L} \left(x_{i,j+j'-1} w_{i,L-j'+1}\right) \qquad (9)
\end{aligned}
$$

Replacing $\mathbf{W}$ with its flipped version $\mathbf{W}'$ which has $w'_{i,j} := w_{i,L-j+1}$, we have

$$
\begin{aligned}
=&\sum_{i=1}^{4} \sum_{j'=1}^{L} \left(x_{i,j+j'-1} w'_{i,j'}\right) \qquad (10) \\
=&\sum_{j'=1}^{L} \left(w'_{f(j+j'-1|\mathbf{X}),j'}\right)
\end{aligned}
$$

Replacing $\mathbf{W}'$ with its exponentiated version $\mathbf{C} := b^{\mathbf{W}'}$, we have

$$
=\sum_{j'=1}^{L} \log_b \left(c_{f(j+j'-1|\mathbf{X}),j'}\right) \qquad (11)
$$

Replacing $\mathbf{C}$ with $\mathbf{P}(\mathbf{W}, b)$ where $(\mathbf{P}(\mathbf{W}, b))_{i,j} := \frac{c_{i,j}}{\sum_{i'=1}^{4} c_{i',j}}$, we have

$$
\begin{aligned}
=&\sum_{j'=1}^{L} \log_b \left( (\mathbf{P}(\mathbf{W}, b))_{f(j+j'-1|\mathbf{X}),j'} \sum_{i=1}^{4} c_{i',j'} \right) \qquad (12) \\
=&\sum_{j'=1}^{L} \log_b \left( (\mathbf{P}(\mathbf{W}, b))_{f(j+j'-1|\mathbf{X}),j'} \right) + \sum_{j'=1}^{L} \log_b \left( \sum_{i'=1}^{4} c_{i',j'} \right)
\end{aligned}
$$

Inserting the probability formula for PWM derived above gives us

$$
=\log_b \text{Prob}(\mathbf{X}[1:4, j:(j+L-1)]|\mathbf{P}(\mathbf{W}, b)) + \sum_{j'=1}^{L} \log_b \left( \sum_{i'=1}^{4} c_{i',j'} \right) \qquad (13)
$$

Noting that the latter term depends entirely on $\mathbf{W}$ and $b$, we have

$$
=\log_b \text{Prob}(\mathbf{X}[1:4, j:(j+L-1)]|\mathbf{P}(\mathbf{W}, b)) + d(\mathbf{W}, b) \qquad (14)
$$

which concludes the proof. □

#### 3 Theorem 2 and its proof

**Theorem 2.** *Assume that the given deep learning framework has its output function* $g(\mathbf{X}|\Theta)$ *(i.e. parameterized by the parameter set* $\Theta$*) of the following form:*

$$g(\mathbf{X}|\Theta) := u\left(h_1\left(\mathbf{X} * \mathbf{W_1}|\theta_1\right), h_2\left(\mathbf{X} * \mathbf{W_2}|\theta_2\right), \cdots, h_k\left(\mathbf{X} * \mathbf{W_k}|\theta_k\right)|\gamma\right) \tag{15}$$

*where* $\mathbf{W_i}, \forall i = 1, \cdots, k$ *are the kernels (and thus part of the parameter set of this model) used for convolution, each* $h_i(\cdot|\theta_i), \forall i = 1, \cdots, k$ *is a function of one of the following forms parameterized by* $\theta_i$*:*

**linear**

$$h_i(\mathbf{x}|\theta_i) = \mathbf{A}_{\theta_i}\mathbf{x} + \mathbf{b}_{\theta_i} \tag{16}$$

**max-linear**

$$h_i(\mathbf{x}|\theta_i) = a_{\theta_i} \max \mathbf{x} + b_{\theta_i} \tag{17}$$

**thresholding-max-linear**

$$h_i(\mathbf{x}|\theta_i) = a_{\theta_i} \max\left(\text{threshold}(\mathbf{x}, z_{\theta_i})\right) + b_{\theta_i} \tag{18}$$

*, and* $u(\cdot|\gamma)$ *an arbitrary function whose paramter is* $\gamma$*. Then for each parameter set* $\Theta^*$*, there are uncountably infinite parameter sets each of which (denoted as* $\Theta'$*) satisfying* $g(\mathbf{X}|\Theta') = g(\mathbf{X}|\Theta^*)$ *when* $\mathbf{X}$ *is fixed.*

*Proof.* The idea of this proof is: for each scaling/shifting of kernels, we could always find a way to modify the parameters of $h_i(\mathbf{x}|\theta_i)$ (i.e. $\theta_i$) to make the output of $h_i(\cdot)$ unchanged.

Assume all kernels are of length $L$ and the input sequence $X$ is of length $N$. Write $\Theta^*$ as $\Theta^* = \{\mathbf{W_1^*}, \cdots, \mathbf{W_k^*}, \theta_1^*, \cdots, \theta_k^*, \gamma^*\}$. Then, due to the identities listed below, $\forall r \in \mathbb{R}^+, t \in \mathbb{R}$, we could always construct $\theta_i'(r,t)$ such that $h_i\left(\mathbf{X} * \mathbf{W_i^*}|\theta_i^*\right) = h_i\left((\mathbf{X} * (r\mathbf{W_i^*} + t))|\theta_i'(r,t)\right)$.

**linear**

$$\begin{aligned}
&h_i\left(\mathbf{X} * \mathbf{W_i^*}|\theta_i^*\right) \\
=&\mathbf{A}_{\theta_i}(\mathbf{X} * \mathbf{W_i^*}) + \mathbf{b}_{\theta_i} \\
=&\frac{1}{r}\mathbf{A}_{\theta_i}(\mathbf{X} * (r\mathbf{W_i^*} + t)) + (\mathbf{b}_{\theta_i} - \frac{tL}{r}\mathbf{A}_{\theta_i}\mathbf{1}_{N-L+1})
\end{aligned} \tag{19}$$

**max-linear**

$$
\begin{aligned}
&h_i\left(\mathbf{X} \ast \mathbf{W_i^*} | \theta_i^*\right) \\
=&a_{\theta_i} \max\left(\mathbf{X} \ast \mathbf{W_i^*}\right) + b_{\theta_i} \\
=&\frac{a_{\theta_i}}{r} \max\left(\mathbf{X} \ast (r\mathbf{W_i^*} + t)\right) + (b_{\theta_i} - \frac{ta_{\theta_i}}{r})
\end{aligned} \tag{20}
$$

**thresholding-max-linear**

$$
\begin{aligned}
&h_i\left(\mathbf{X} \ast \mathbf{W_i^*} | \theta_i^*\right) \\
=&a_{\theta_i} \max\left(\text{threshold}(\mathbf{X} \ast \mathbf{W_i^*}, z_{\theta_i})\right) + b_{\theta_i} \\
=&\frac{a_{\theta_i}}{r} \max\left(\text{threshold}(\mathbf{X} \ast (r\mathbf{W_i^*} + t), rz_{\theta_i} + t)\right) + (b_{\theta_i} - \frac{ta_{\theta_i}}{r})
\end{aligned} \tag{21}
$$

Specifically, the construction of $\theta_i'(r, t)$ is:

**linear**

$$
\mathbf{A}_{\theta_i'} := \frac{1}{r}\mathbf{A}_{\theta_i}, \mathbf{b}_{\theta_i'} := (\mathbf{b}_{\theta_i} - \frac{t}{r}\mathbf{A}_{\theta_i}) \tag{22}
$$

**max-linear**

$$
a_{\theta_i'} := \frac{a_{\theta_i}}{r}, b_{\theta_i'} := (b_{\theta_i} - \frac{ta_{\theta_i}}{r}) \tag{23}
$$

**thresholding-max-linear**

$$
z_{\theta_i'} := rz_{\theta_i} + t, a_{\theta_i'} := \frac{a_{\theta_i}}{r}, b_{\theta_i'} := (b_{\theta_i} - \frac{ta_{\theta_i}}{r}) \tag{24}
$$

Therefore, if we define $\mathbf{W_i'} := r\mathbf{W_i^*}$, then $\Theta_i' := \{\mathbf{W_1'}, \cdots, \mathbf{W_k'}, \theta_1', \cdots, \theta_k', \gamma^*\}$ satisfies the identity $g(\mathbf{X}|\Theta') = g(\mathbf{X}|\Theta^*)$.

This implies that for each solution and each pair of $(r, t)$ that changes the i-th kernel $\mathbf{W_i}$, we could always find some parameters for $h_i(\cdot)$ to build up a new $\Theta_i'$ that keep the output of $h_i(\cdot)$, and thus the output of $g(\cdot)$, unchanged. Since the number of all possible pairs of $(r, t)$ (including the case $(r, t = 0)$) is uncountably infinite, we could create uncountably infinite number of such $\Theta_i'$.

□

When all $h_i(\cdot)$ are thresholding-max-linear with $z_{\theta_i} = 0$, $g(\cdot)$ is essentially a convolution layer with ReLU activation, followed by a global max-pooling layer, then by other arbitrary layers. If the thresholds $(z_{\theta_i})$ are allowed to change, then

Theorem 2 could be applied directly; if not, Theorem 2 could still be applied by enforcing that $t$ must be 0 no matter what $r$ is chosen.

Also, note that other activation functions and other model structures might also make Theorem 2 hold, as long as a similar construction is given. For example,

1. A global average pooling could be linear by requiring that the input sequences have the same length, and thus Theorem 2 could be applied to a CNN framework where convolution is followed by a global average pooling.

2. A local max-pooling $h(\mathbf{c})$ is equivalent to the concatenation of the output of a series of $h_i(\mathbf{c}) := \max(l(\mathbf{c}))$ , where each $h_i(\mathbf{c})$ is applied to a separate window from the local pooling, and $l(\mathbf{c})$ is a linear transformation that adds a very negative value to all elements in $\mathbf{c}$ out of the window, and 0 to elements in the window. Since Theorem 2 ensures that each $h_i(\mathbf{c})$ could have their output unchanged regardless of how the kernel changes, it could also be applied to $h(\mathbf{c})$.

3. Similarly, Theorem 2 also holds for local average-pooling.

4. Since the $h(\cdot)$'s are independent of each other, Theorem 2 also holds for CNN frameworks where multiple layers are connected to the convolution output at the same time (e.g., by connecting to the convolution a global max-pooling layer and also a global average-pooling layer).

Therefore, Theorem 2 could be accessed by a variety of CNN frameworks that have been used for handling DNA/RNA sequences (e.g., [1–3] ).

### 4 Corollaries 1 and 2 and their deduction

**Corollary 1.** *For each sequence profile, there are uncountably infinite other sequence profiles that classify sequences in the same way as this sequence profile does when used in deep learning frameworks with output function $g(\cdot)$ as described in Theorem 2.*

*Proof.* In the following deduction, only models whose structure satisfying those criteria in Theorem 2 will be considered. In addition, we will only prove the special case where there is only one kernel in the convolution layer applied to the input sequences directly, as the proof could obviously extends to models with multiple kernels.

According to the transformation the resulting PWM $\mathbf{P}(\mathbf{W}, b)$ satisfies $(\mathbf{P}(\mathbf{W}, b))_{i,j'} = \frac{c_{i,j'}}{\sum_{i'=1}^{4} c_{i',j'}} = \frac{b^{w'_{i,j'}}}{\sum_{i'=1}^{4} b^{w'_{i',j'}}} = \frac{b^{w_{i,L-j'+1}}}{\sum_{i'=1}^{4} b^{w_{i',L-j'+1}}}$.

As for its performance on sequence regression/classification, note that for any kernel $\mathbf{W}$, we have the identity in Theorem 1

$$\begin{aligned}(\mathbf{X} \ast \mathbf{W})_j &= \sum_{j'=1}^{L} \log_b\left((\mathbf{P}(\mathbf{W},b))_{f(j+j'-1|\mathbf{X}),j'}\right) + \sum_{j'=1}^{L} \log_b\left(\sum_{i'=1}^{4} c_{i',j'}\right) \\ &= \sum_{j'=1}^{L} \log_b\left((\mathbf{P}(\mathbf{W},b))_{f(j+j'-1|\mathbf{X}),j'}\right) + \sum_{j'=1}^{L} \log_b\left(\sum_{i'=1}^{4} b^{w'_{i',j'}}\right) \\ &= \sum_{j'=1}^{L} \log_b\left((\mathbf{P}(\mathbf{W},b))_{f(j+j'-1|\mathbf{X}),j'}\right) + \sum_{j'=1}^{L} \log_b\left(\sum_{i'=1}^{4} b^{w_{i',L-j'+1}}\right) \\ &= \sum_{j'=1}^{L} \log_b\left((\mathbf{P}(\mathbf{W},b))_{f(j+j'-1|\mathbf{X}),j'}\right) + \sum_{j'=1}^{L} \log_b\left(\sum_{i'=1}^{4} b^{w_{i',j'}}\right)\end{aligned} \tag{25}$$

Taking advantage of the one-hot encoding nature of $\mathbf{X}$, we have

$$(\mathbf{X} \ast (\mathbf{W} - \frac{1}{L}\sum_{j'=1}^{L} \log_b\left(\sum_{i'=1}^{4} b^{w_{i',j'}}\right)))_j = \sum_{j'=1}^{L} \log_b\left((\mathbf{P}(\mathbf{W},b))_{f(j+j'-1|\mathbf{X}),j'}\right) \tag{26}$$

By Theorem 2, $\mathbf{W}$ and $(\mathbf{W} - \frac{1}{L}\sum_{j'=1}^{L} \log_b\left(\sum_{i'=1}^{4} b^{w_{i',j'}}\right))$ (when regarded as a new kernel) could regress/classify sequences exactly the same. Since the latter is identical to the log-likelihood computed by $\mathbf{P}(\mathbf{W},b)$, we could say that $\mathbf{W}$ and $\mathbf{P}(\mathbf{W},b)$ could also regress/classify sequences exactly the same.

Now let's check what will happen to the PWM itself and its performance on sequence regression/classification, if the base for calculating the log-likelihood is changed.

Without loss of generality, if we replace $b$ with $b^r (r > 0, r \neq 1)$, the new PWM $\mathbf{P}(\mathbf{W},b^r)$ satisfies $(\mathbf{P}(\mathbf{W},b^r))_{i,j'} = \frac{(b^r)^{w_{i,L-j'+1}}}{\sum_{i'=1}^{4}(b^r)^{w_{i',L-j'+1}}}$. Then it can be shown that changing the base will definitely change the resulting PWM, unless each column of the kernel is filled with identical elements (which is rarely possible). Specifically, suppose there is some $r > 0, r \neq 1$ that will not change the resulting PWM for some kernel $\mathbf{W}$:

$$(\mathbf{P}(\mathbf{W},b))_{i,j'} = (\mathbf{P}(\mathbf{W},b^r))_{i,j'} \tag{27}$$

i.e.,

$$\frac{b^{w_{i,L-j'+1}}}{\sum_{i'=1}^{4} b^{w_{i',L-j'+1}}} = \frac{(b^r)^{w_{i,L-j'+1}}}{\sum_{i'=1}^{4}(b^r)^{w_{i',L-j'+1}}} \tag{28}$$

then we have

$$\frac{b^{w_{i,L-j'+1}}}{(b^r)^{w_{i,L-j'+1}}} = \frac{\sum_{i'=1}^{4} b^{w_{i',L-j'+1}}}{\sum_{i'=1}^{4} (b^r)^{w_{i',L-j'+1}}} \tag{29}$$

since the RHS does not depend on $i$, we can, for example, replace $i$ with 1 and again with 2, and have

$$\frac{b^{w_{1,L-j'+1}}}{(b^r)^{w_{1,L-j'+1}}} = \frac{b^{w_{2,L-j'+1}}}{(b^r)^{w_{2,L-j'+1}}} \tag{30}$$

therefore

$$\frac{b^{w_{1,L-j'+1}}}{b^{w_{2,L-j'+1}}} = \frac{(b^r)^{w_{1,L-j'+1}}}{(b^r)^{w_{2,L-j'+1}}} = \left(\frac{b^{w_{1,L-j'+1}}}{b^{w_{2,L-j'+1}}}\right)^r \tag{31}$$

because $b > 0$, $\frac{b^{w_{1,L-j'+1}}}{b^{w_{2,L-j'+1}}} > 0$, and we have

$$1 = \left(\frac{b^{w_{1,L-j'+1}}}{b^{w_{2,L-j'+1}}}\right)^{r-1} \tag{32}$$

now we use the assumption that $r - 1 \neq 0$, and because $\frac{b^{w_{1,L-j'+1}}}{b^{w_{2,L-j'+1}}} > 0$, we must have

$$\frac{b^{w_{1,L-j'+1}}}{b^{w_{2,L-j'+1}}} = 1 \tag{33}$$

i.e.,

$$w_{1,L-j'+1} = w_{2,L-j'+1} \tag{34}$$

but note that the deduction above does not depend on specific values of $i$, and thus we have

$$w_{1,L-j'+1} = w_{2,L-j'+1} = w_{3,L-j'+1} = w_{4,L-j'+1} \tag{35}$$

which concludes the proof. Note that when $r \neq 1$ but changing the base does not change the resulting PWM, the resulting PWM must be a matrix filled with 0.25. Such kernel (and the resulting PWM) is impossible to distinguish any two sequences from each other, and thus is very unlikely to be from a trained model that regress/classify sequences well.

However, a change in the resulting PWM is not accompanied with a change in the performance. Note that the identity below:

$$(\mathbf{X} * \mathbf{W})_j = \sum_{j'=1}^{L} \log_{b^r} \left( (\mathbf{P}(\mathbf{W}, b^r))_{f(j+j'-1|\mathbf{X}),j'} \right) + \sum_{j'=1}^{L} \log_{b^r} \left( \sum_{i'=1}^{4} (b^r)^{w_{i',j'}} \right) \quad (36)$$

could be rearranged in a way similar to the previous one:

$$\begin{aligned} &(\mathbf{X} * (r\mathbf{W} - \frac{1}{L} \sum_{j'=1}^{L} \log_b \left( \sum_{i'=1}^{4} (b^r)^{w_{i',j'}} \right)))_j \\ &= \sum_{j'=1}^{L} \log_b \left( (\mathbf{P}(\mathbf{W}, b^r))_{f(j+j'-1|\mathbf{X}),j'} \right) \end{aligned} \quad (37)$$

Again by Theorem 2, $\mathbf{W}$ and $(r\mathbf{W} - \frac{1}{L} \sum_{j'=1}^{L} \log_b \left( \sum_{i'=1}^{4} (b^r)^{w_{i',j'}} \right))$ (when regarded as a new kernel), and thus $\mathbf{P}(\mathbf{W}, b^r)$, could regress/classify sequences exactly the same.

Combining all the equivalence relationships deduced above, we arrived at the conclusion: $\mathbf{P}(\mathbf{W}, b)$ and $\mathbf{P}(\mathbf{W}, b^r)$ could also regress/classify sequences exactly the same. Since $r$ could be any positive real number in $\mathbb{R}$, for each PWM $\mathbf{P}(\mathbf{W}, b)$ we could always construct uncountably infinite $\mathbf{P}(\mathbf{W}, b^r)$'s that has the same performance on regression/classification, thus concluding the proof. □

This raises the concern that additional constraints must be applied when selecting the optimal kernels (and thus sequence profiles) based on the trained model. A popular choice would be to use maximum likelihood estimation to determine the base $b$, which will be introduced in the next section.

On the other hand, it doesn't escape from our attention that this corollary also suggests that the classical sequence logo (e.g., the one used in Deepbind's paper [1] ) does not fit CNN-oriented profile visualization.

**Corollary 2.** $\forall \mathbf{W}^{**}, \mathbf{W}^{*}$*, we have:* $\forall b (b > 1), \mathbf{P}(\mathbf{W}^{**}, b) = \mathbf{P}(\mathbf{W}^{*}, b)$ *if and only if* $\forall i \in \{1, \cdots, 4\}, j \in \{1, \cdots, L\}, w_{i,j}^{**} = w_{i,j}^{*} + t_j$*, where* $t_j \in \mathbb{R}$ *is independent of* $\mathbf{W}^{**}$ *and* $\mathbf{W}^{*}$ *.*

*Proof.* We prove the "if" direction and the "only-if" direction separately.

**The "if" direction** This is a natural result of the transformation. Specifically, we have

$$(\mathbf{P}(\mathbf{W}, b))_{i,j} = \frac{c_{i,j}}{\sum_{i'=1}^{4} c_{i',j}} = \frac{b^{w'_{i,j}}}{\sum_{i'=1}^{4} b^{w'_{i',j}}} = \frac{b^{w_{i,L-j+1}}}{\sum_{i'=1}^{4} b^{w_{i',L-j+1}}} \tag{38}$$

And therefore

$$\begin{aligned} &(\mathbf{P}(\mathbf{W}^{**}, b))_{i,j} \\ =&\frac{b^{w^{**}_{i,L-j+1}}}{\sum_{i'=1}^{4} b^{w^{**}_{i',L-j+1}}} = \frac{b^{w^{*}_{i,L-j+1}+t_j}}{\sum_{i'=1}^{4} b^{w^{*}_{i',L-j+1}+t_j}} = \frac{b^{w^{*}_{i,L-j+1}}}{\sum_{i'=1}^{4} b^{w^{*}_{i',L-j+1}}} \\ =&(\mathbf{P}(\mathbf{W}^{*}, b))_{i,j} \end{aligned} \tag{39}$$

which concludes the proof.

**The "only-if" direction** Fix $j \in \{1, \cdots, L\}$. As in the proof of the "if" direction, we already have for any $\mathbf{W}$

$$(\mathbf{P}(\mathbf{W}, b))_{i,j} = \frac{b^{w_{i,L-j+1}}}{\sum_{i'=1}^{4} b^{w_{i',L-j+1}}} \tag{40}$$

Therefore, we have

$$\begin{aligned} &(\mathbf{P}(\mathbf{W}, b))_{1,j} : (\mathbf{P}(\mathbf{W}, b))_{2,j} : (\mathbf{P}(\mathbf{W}, b))_{3,j} : (\mathbf{P}(\mathbf{W}, b))_{4,j} \\ =&b^{w_{1,L-j+1}} : b^{w_{2,L-j+1}} : b^{w_{3,L-j+1}} : b^{w_{4,L-j+1}} \end{aligned} \tag{41}$$

Now that we have $\mathbf{P}(\mathbf{W}^{**}, b) = \mathbf{P}(\mathbf{W}^{*}, b)$, we could get the following result:

$$\begin{aligned} &b^{w^{**}_{1,L-j+1}} : b^{w^{**}_{2,L-j+1}} : b^{w^{**}_{3,L-j+1}} : b^{w^{**}_{4,L-j+1}} \\ =&b^{w^{*}_{1,L-j+1}} : b^{w^{*}_{2,L-j+1}} : b^{w^{*}_{3,L-j+1}} : b^{w^{*}_{4,L-j+1}} \end{aligned} \tag{42}$$

Therefore, $b^{w^{**}_{i,L-j+1}} = T_j b^{w^{*}_{i,L-j+1}}$ for some positive constant $T_j$ that is independent of $\mathbf{W}^{**}$ and $\mathbf{W}^{*}$. Taking the logarithm with base $b$ on both sides gives $w^{**}_{i,L-j+1} = \log_b T_j + w^{*}_{i,L-j+1}$, which concludes the proof if we set $t_{L-j+1} := \log_b T_j$.

□

This is useful if one would like to make some predefined PWM (with the restriction that no 0's or 1's are present) work like kernels in a popular type of CNN model where the thresholding-max-linear structure is used with the threshold being 0 (i.e. ReLU activation is used), without changing its capability of regressing/classifying sequences.

1. Generally, if we transform a PWM back to the kernel by reversing the transformation described in the main text, we could use this corollary (specifically, the "only-if" direction) to assume that the constant difference between convolution and log-likelihood (i.e. $\sum_{j'=1}^{L} \log_b \left( \sum_{i=1}^{4} (b)^{w_{i,j'}} \right)$) is 0 without worrying about changing the underlying PWM; this allows us to skip the reversed normalization step (i.e. $\mathbf{P}(\mathbf{W}, b) \mapsto \mathbf{C}$); the rest two steps (i.e. $\mathbf{C} \mapsto \mathbf{W}', \mathbf{W}' \mapsto \mathbf{W}$ ) are both invertible and could be carried out unambiguously.

2. Since all elements in $\mathbf{P}(\mathbf{W}, b)$ are within $(0, 1)$, we would end up with a kernel filled with negative elements only. Such negativeness will not be a problem if the CNN model has a linear activation, but it will be if the activation is ReLU: the convolution would be a constant of 0, thus wiping out any potential signals. We could, however, use this corollary (the "if" direction) again to avoid this problem without (again) worrying about changing the the underlying PWM: just apply to each of the kernels a positive shift (which could be predefined or be trained) that is big enough to recover the signals of interest from input sequences.

3. In fact, this problem could be completely circumvented, and we could make the kernels classify sequences as good as the original PWMs. Specifically, for each kernel we could make the absolute value of the shift so big that it is impossible to find a sequence whose maximum of convolution by the kernel is negative (which is easy to implement by using as the shift, for example, the absolute value of the smallest element in the kernel); in this case, the ReLU activation behaves like the linear activation precisely and, due to Theorem 2, for the sequence classification problem the shifted kernels under ReLU activation could perform as good as the unshifted kernels perform under linear activation, and thus as good as the original PWMs perform (using log-likelihood).

### 5 Selecting the best PWM for a given kernel by maximum likelihood estimation

Here we establish a maximum likelihood estimation (MLE) of optimal PWM sets (specifically, the optimal $b$) for CNN models with the popular structure "input → convolution → activation → global max-pooling → arbitrary layers", where the activation could be either linear or ReLU, or some other function that is possible to preserve prediction performance in Theorem 2. The MLE for CNN models with other structures could be derived in a similar manner.

Since the activation is monotonically increasing, the output of the model will not change if we switch the activation with the global max-pooling layer. In this way, the model becomes "input → convolution → global max-pooling → activation → arbitrary layers", and a reasonable (and simple) likelihood could be the joint probability of observing all max-scored fragments given their generative PWMs transformed from all the kernels with base $b$. Of course, if biologically plausible one could argue further (and thus modify the likelihood accordingly) that for a certain kernel, sequences whose convolution by that kernel do not pass the activation should be excluded from the joint probability.

Recall that from Theorem 1 we have

$$(\mathbf{X} * \mathbf{W})_j = \log_b \mathrm{Prob}(\mathbf{X}[1:4, j:(j+L-1)]|\mathbf{P}(\mathbf{W}, b)) + d(\mathbf{W}, b) \tag{43}$$

where $d(\mathbf{W}, b) = \sum_{j'=1}^{L} \log_b \left(\sum_{i'=1}^{4} c_{i',j'}\right) = \sum_{j'=1}^{L} \log_b \left(\sum_{i'=1}^{4} b^{w_{i',j'}}\right)$.

Now assume that we have $n$ input sequences, $\mathbf{X}^{(1)}, \cdots, \mathbf{X}^{(n)}$, and $k$ kernels, $\mathbf{W}^{(1)}, \cdots, \mathbf{W}^{(k)}$. Define $j^*(s,t) := \arg\max_j \mathrm{Prob}(\mathbf{X}^{(s)}[1:4, j:(j+L-1)]|\mathbf{P}(\mathbf{W}^{(t)}, b))$, i.e. the start coordinate of the fragment from $\mathbf{X}^{(s)}$ that is the most possible one to be generated from $\mathbf{P}(\mathbf{W}^{(t)}, b)$. Note that $j^*(s,t)$ does not change when all quantities but $b$ do not change, as $j^*(s,t)$ could be derived from the $b$-independent convolution $\mathbf{X}^{(s)} * \mathbf{W}^{(t)}$; therefore, we could treat $j^*(s,t)$ as constants when finding the optimal $b$.

We then could write the following likelihood $q(\mathbf{X}^{(1)}, \cdots, \mathbf{X}^{(n)}|\mathbf{W}^{(1)}, \cdots, \mathbf{W}^{(k)}, b)$:

$$\begin{aligned} & q(\mathbf{X}^{(1)}, \cdots, \mathbf{X}^{(n)}|\mathbf{W}^{(1)}, \cdots, \mathbf{W}^{(k)}, b) \\ = & \prod_{s=1}^{n}\prod_{t=1}^{k} \mathrm{Prob}(\mathbf{X}^{(s)}[1:4, j^*(s,t):(j^*(s,t)+L-1)]|\mathbf{P}(\mathbf{W}^{(t)}, b)) \end{aligned} \tag{44}$$

$$=\prod_{s=1}^{n}\prod_{t=1}^{k} e^{(\ln b)\log_b \text{Prob}(\mathbf{X}^{(s)}[1:4,j*(s,t):(j*(s,t)+L-1)]|\mathbf{P}(\mathbf{W}^{(t)},b))}$$

$$=\prod_{s=1}^{n}\prod_{t=1}^{k} e^{(\ln b)(\mathbf{X}^{(s)}*\mathbf{W}^{(t)})_{j*(s,t)}-(\ln b)\sum_{j'=1}^{L}\log_b\left(\sum_{i'=1}^{4} b^{w_{i',j'}^{(t)}}\right)}$$

Taking the natural logarithm of both sides gives us

$$\begin{aligned}
&\ln q(\mathbf{X}^{(1)},\cdots,\mathbf{X}^{(n)}|\mathbf{W}^{(1)},\cdots,\mathbf{W}^{(k)}) \\
=&\sum_{s=1}^{n}\sum_{t=1}^{k}\left((\ln b)(\mathbf{X}^{(s)}*\mathbf{W}^{(t)})_{j*(s,t)}-\sum_{j'=1}^{L}\ln\left(\sum_{i'=1}^{4} b^{w_{i',j'}^{(t)}}\right)\right) \qquad (45)\\
=&(\ln b)\sum_{s=1}^{n}\sum_{t=1}^{k}(\mathbf{X}^{(s)}*\mathbf{W}^{(t)})_{j*(s,t)}-n\sum_{t=1}^{k}\sum_{j'=1}^{L}\ln\left(\sum_{i'=1}^{4} e^{(\ln b)w_{i',j'}^{(t)}}\right)
\end{aligned}$$

Therefore, we have the following maximum likelihood estimation:

$$\begin{aligned}
&\arg\max_b q(\mathbf{X}^{(1)},\cdots,\mathbf{X}^{(n)}|\mathbf{W}^{(1)},\cdots,\mathbf{W}^{(k)},b) \\
=&\arg\max_b \ln q(\mathbf{X}^{(1)},\cdots,\mathbf{X}^{(n)}|\mathbf{W}^{(1)},\cdots,\mathbf{W}^{(k)},b) \qquad (46)\\
=&\arg\max_b\left((\ln b)\sum_{s=1}^{n}\sum_{t=1}^{k}(\mathbf{X}^{(s)}*\mathbf{W}^{(t)})_{j*(s,t)}-n\sum_{t=1}^{k}\sum_{j'=1}^{L}\ln\left(\sum_{i'=1}^{4} e^{(\ln b)w_{i',j'}^{(t)}}\right)\right)\\
=&\arg\max_{v:=\ln b}\left(v\sum_{s=1}^{n}\sum_{t=1}^{k}(\mathbf{X}^{(s)}*\mathbf{W}^{(t)})_{j*(s,t)}-n\sum_{t=1}^{k}\sum_{j'=1}^{L}\ln\left(\sum_{i'=1}^{4} e^{vw_{i',j'}^{(t)}}\right)\right)
\end{aligned}$$

Although the convolution itself does not involve $b$ (or $v$), the likelihood function has a form of logarithm of sum of exponentials of $b$ (or $v$), which makes it very hard to derive a closed-form solution. In fact, even evaluating the first derivative ($\frac{dq}{db}$ or $\frac{dq}{dv}$) is very cumbersome, and therefore a derivative-free optimizer might be easier to use here.

### 6 Comments on the special case where the sequence contains N

Sometimes the input sequences contain a special type of "nucleotide", N, due to one of the following reasons:

- Some of the nucleotides in the input sequence are inherently unknown due to various reasons (e.g., low quality of sequencing).
- The input sequences are of equal length and must be padded before, after, or around with N.
- Certain models assume that the input sequence must be concatenated from several sequence fragments, in a way that scanning one such fragment will not affect scanning others. Therefore, these fragments are concatenated by a stretch of N's (typically not shorter than the length of kernels).

In these cases, N (on position $j$ ) is generally encoded by one of the following two ways:

1. By zeros: $x_{i,j} := 0$ for $i = 1, 2, 3, 4$
2. By one quarters (0.25): $x_{i,j} := 0.25$ for $i = 1, 2, 3, 4$

In either cases, Theorem 1 still holds, except that the constant term $d(\mathbf{W}, b)$ now depends further on the positions of N's. Formally, we have:

**Theorem 3.** *$\forall b > 1$ and $\forall \mathbf{X}$ with $N$ columns, where the position set $J (0 \leqslant J < N)$, i.e., $j_1, j_2, \cdots, j_{|J|} (j_1 < j_2 < \cdots < j_{|J|})$, are nucleotides of A, C, G, or T, and other $N - J$ positions are nucleotides of N. Then the convolution of $\mathbf{X}$ with a kernel $\mathbf{W}$ of length $L$ has:*

$$(\mathbf{X} * \mathbf{W})_j \tag{47}$$

$$\begin{aligned}
&= \log_b \mathrm{Prob}(\mathbf{X}[1:4, \mathit{order}(J'(j))] | \mathbf{P}(\mathbf{W}[1:4, \mathit{order}(\{L - (j' - j) + 1 | j' \in J'(j)\})], b)) \\
&\quad + d(\mathbf{W}[1:4, \mathit{order}(\{L - (j' - j) + 1 | j' \in J'(j)\})], b) \\
&\quad + \alpha \times \sum_{i=1}^{4} \sum_{j^@=1}^{|J^*(j)|} (\log_b \mathbf{P}(\mathbf{W}[1:4, \mathit{order}(\{L - (j^* - j) + 1 | j^* \in J^*(j)\})], b))_{i, j^@} \\
&\quad + \beta \times d(\mathbf{W}[1:4, \mathit{order}(\{L - (j^* - j) + 1 | j^* \in J^*(j)\})], b)
\end{aligned} \tag{48}$$

*where*

1. *$J'(j) = J \cap \{i \in \mathbb{Z} | j \leqslant i \leqslant j + L - 1\}$ is the set of all non-N positions within the window $j : (j + L - 1)$ of $\mathbf{X}$;*
2. *$J^*(j) = \{i \in \mathbb{Z} | j \leqslant i \leqslant j + L - 1\} - J'(j)$ is the set of all N positions in this window; and*

*3. d follows the definition in Theorem 1.*

*4. If N's are encoded by 0, then $\alpha = \beta = 0$; if by 0.25, then $\alpha = 0.25, \beta = 1$.*

Note that we deliberately omit the case where the sequence consists entirely of N, because such sequence contains no signal and thus is not meaningful to scan.

Basically, this theorem states that the calculation of the log-likelihood itself does not take into account the positions of N (the PWM does not consider these positions), and the constant depends on which positions are N in the window of question, and on how the N is encoded.

*Proof.* Let's first split the window $j : (j + L - 1)$ of $\mathbf{X}$ into two parts:

1. $\mathbf{X}[1 : 4, order(J'(j))]$, the one with all non-N positions.

2. $\mathbf{X}[1 : 4, order(J^*(j))]$, the one with all N positions.

By Theorem 1, and noting that the position $j'$ in $\mathbf{X}$ maps in convolution to the position $L - (j' - j) + 1$ in $\mathbf{W}$, for the first part we have

$$
\begin{aligned}
&(\mathbf{X}[1 : 4, order(J'(j))] \circledast \mathbf{W})_j \\
=& \log_b \text{Prob}(\mathbf{X}[1 : 4, order(J'(j))] | \mathbf{P}(\mathbf{W}[1 : 4, order(\{L - (j' - j) + 1 | j' \in J'(j)\})], b)) \\
&+ d(\mathbf{W}[1 : 4, order(\{L - (j' - j) + 1 | j' \in J'(j)\})], b) \qquad (49)
\end{aligned}
$$

For the second part,

1. If N's are encoded by 0, then their convolution is 0.

2. If N's are encoded by 0.25, then their convolution is essentially an average of the following four sequences of equal length: a string of A's, a string of C's, a string of G's, and a string of T's. By Theorem 1, each of these sequence (denoted by $\mathbf{B}$) has the following :

$$
\begin{aligned}
&(\mathbf{B} \circledast \mathbf{W}[1 : 4, order(\{L - (j^* - j) + 1 | j^* \in J^*(j)\})])_j \\
=& \log_b \text{Prob}(\mathbf{B} | \mathbf{P}(\mathbf{W}[1 : 4, order(\{L - (j' - j) + 1 | j^* \in J^*(j)\})], b)) \\
&+ d(\mathbf{W}[1 : 4, order(\{L - (j^* - j) + 1 | j^* \in J^*(j)\})], b) \qquad (50)
\end{aligned}
$$

During the average, the $log_b$ items on RHS cover all elements in $\mathbf{P}(\mathbf{W}[1 : 4, order(\{L - (j' - j) + 1 | j^* \in J^*(j)\})], b)$ and is equal to:

$0.25 \sum_{i=1}^{4} \sum_{j^@=1}^{|J^*(j)|} \log_b (\mathbf{P}(\mathbf{W}[1 : 4, order(\{L - (j^* - j) + 1 | j^* \in J^*(j)\})], b))_{i,j^@}$.

The $d$ items are the same and thus unchanged.

Adding the two parts above gives the theorem. □
