## Supplementary Table 1 for "An exact transformation for CNN kernel enables accurate sequence motif identification and leads to a potentially full probabilistic interpretation of CNN"

Supplementary Table 1. Analysis setup for simulation

| Motif type | Sequence-conserved | Structure-conserved |
| --- | --- | --- |
| Source | JASPAR CORE 2016 (Mathelier *et al.*, 2016), 1082 motifs | Rfam version 12.2 (Nawrocki *et al.*, 2015), 2588 motifs |
| Count of positive sequences | 1500 | |
| Count of negative sequences | 1500 | |
| Sequence length | 50 | 5000 |
| Generation of each positive sequence | Firstly, sample a fragment from the motif’s PWM.  Then extend the fragment from both sides with random sequences of random lengths. Each nucleotide in each random sequence is sampled from an independent uniform distribution (Prob(A) = Prob(C) = Prob(G) = Prob(T) = 0.25). | Sample a sequence using cmemit from Infernal version 1.1.2 (Nawrocki and Eddy, 2013), with -l set |
| Generation of each negative sequence | Generate a random sequence, in which each nucleotide is sampled from an independent uniform distribution (Prob(A) = Prob(C) = Prob(G) = Prob(T) = 0.25). | Sample a sequence using cmemit from Infernal version 1.1.2 (Nawrocki and Eddy, 2013), with -l set and --exp set to 0.01 |
| Encoding of the sequence into input | One-hot encoding (See Definition, Common 3 in Supplementary Notes for its definition) | |
| CNN structure | Input -> Convolution -> ReLU activation -> Global Maxpooling -> Logistic regression -> Output, modelled by keras (Chollet and others, 2015) with the theano backend (The Theano Development Team *et al.*, 2016). | |
| CNN kernel count | 1 | 5 |
| CNN kernel length | Same as the length of the motif to detect | 80 |
| Training strategy | 80% of the dataset is for training and 20% is for validation.  Loss function is binary cross-entropy.  Optimizer is RMSProp with default parameters in keras (Chollet and others, 2015) (learning rate = 0.001, rho = 0.9, epsilon = 1e-08).  Batch size is set to 100.  Stop training when at least one of the following two conditions is met: [1] validation loss does not increase after 6 epochs (the initial loss is set to infinity); [2] 60 epochs have been finished. | |
| Details of the exactPWM transformation | e (the base of natural logarithm) was used as the base b. | |
| Details of the heuristic PWM transformation | The implementation follows the weighted transformation described in (Alipanahi *et al.*, 2015). All sequences from the training and the validation datasets were used. | |
| Prediction for a single input sequence based on log-likelihood of transformed PWMs (for AUC computation) | The log-likelihood of the PWM of this sequence.  For a given motif, if the resulting AUC was less than 0.5, however, the log-likelihoods were multiplied by -1 and used as the prediction instead. | The probabilistic prediction for that sequence from a logistic regression model, which was trained with all PWMs’ log-likelihoods as inputs and with the sequence label as output. Logistic regression was trained using sckit-learn (Pedregosa *et al.*, 2011). |
