## Supplementary Table 2 for "An exact transformation for CNN kernel enables accurate sequence motif identification and leads to a potentially full probabilistic interpretation of CNN"

Supplementary Table 2. Analysis setup for Figure 3

| Source of dataset | Deepbind’s dataset (downloaded from http://tools.genes.toronto.edu/deepbind/nbtcode/nbt3300-supplementary-software.zip) |
| --- | --- |
| Source of model structure and parameters | Structure was constructed as specified by Deepbind itself (Alipanahi *et al.*, 2015).  Parameters, including hyperparameters (e.g., the number of kernels), were from Deepbind’s specification file (downloaded from <http://tools.genes.toronto.edu/deepbind/nbtcode/nbt3300-supplementary-software.zip)>.  Generally, for motifs we selected their Deepbind models have the following structure:  1. The input sequence is fed to a convolutional layer with a series of kernels and ReLU as the activation function.  2. The output of this convolutional layer is fed to a global max-pooling layer.  3. The input sequence is reverse-complemented, and fed to the same convolutional layer and then a global max-pooling layer.  4. Now each kernel has two pooling results for each input sequence, one from the sequence itself and the other from its reverse complement. The larger one was kept.  5. The resulting sample matrix ( the larger pooling result of each kernel for each input sequence) is then fed to either a linear regression directly, or a neural network with one hidden layer and ReLU activation. |
| Choice of benchmarking dataset | Each model has two different datasets, one named with “A” or “AC”, and the other with “B”. We chose the one with “A” or “AC” as the benchmarking dataset.  To simplify the downstream analysis, we also filtered for sequences that pass ReLU activation for all kernels. |
| Encoding of the sequence into input | One-hot encoding (See Definition, Common 3 in Supplementary Notes for its definition) |
| Sequence preprocessing | Each input sequence was padded with 0.25 on BOTH sides. The padding length on each side was equal to the length of convolutional kernel minus 1. In addition, sequences containing the character ‘N’ or ‘n’ were discarded. |
| CNN implementation | Modelled by keras (Chollet and others, 2015) with the theano backend (The Theano Development Team *et al.*, 2016). |
| Details of the exact PWM transformation | e^5^ was used as the base of logarithm b. We have attempted to use MLE to estimate the optimal base of logarithm, but it resulted in numerical overflow for certain kernels; therefore, we gave up using MLE and switched to a moderate base of logarithm. |
| Details of the heuristic PWM transformation | The implementation follows the transformation described in (Alipanahi *et al.*, 2015), with the exception that for each subsequence index that goes beyond the extent of “s” (the test sequence), we added the padded value (0.25) to A, C, G, and T for that position instead of adopting the special ‘empty’ character. We noted that, as specified in (Alipanahi *et al.*, 2015), only sequences passing the ReLU activation were considered.  All sequences from the benchmarking dataset were used. |
| Special treatment to the calculation of log-likelihoods for 0.25-padded sequences | The padded regions were also considered during the calculation of log-likelihoods by summing all log-probabilities within such regions, multiplying them with 0.25, and adding them to the log-likelihood calculated on the original sequence. See Theorem 4 in the Supplemental Notes for more details. |
| Special treatment to the calculation of log-likelihoods for kernels from CNN models supporting reverse complements | Because the original CNN model takes into account the reverse complement of the input sequence, the log-likelihood we calculated here must also take it into account. Specifically, for each PWM and each sequence, the log-likelihood was calculated as follows:  1. We calculated the maximal log-likelihood of that PWM on that sequence;  2. We calculated the maximal log-likelihood of that PWM on the reverse complement of that sequence;  3. We took the larger one of the two above as the final log-likelihood of that PWM on that sequence.  Here the base of the logarithm is the base of the natural logarithm. |
| Prediction for a single input sequence based on log-likelihood of transformed PWMs (for MAPE and MSE computation) | A model was constructed with the same hyperparameter setting as that in the last part of “Source of model structure and parameters” (i.e., either a linear regression or a neural network with one hidden layer and ReLU activation).  For each sequence, the input to this model was the final log-likelihoods of transformed PWMs on that sequence.  The expected output of this model was the sequence’s output in the benchmarking dataset.  The benchmarking dataset was randomly split into training, validation, and testing datasets with sample ratio 0.60:0.15:0.25.  For the derivation of model parameters for PWMs transformed by the exact transformation, see the next row of table.  For the re-training of models for PWMs transformed by the heuristic transformation:  The model was trained with loss being MSE, optimizer being RMSprop (learning rate = 0.001, rho = 0.9, epsilon = 1e-08), and batch size being 100. The training stopped once one of the two conditions was met: [1] validation loss does not increase after 21 epochs (the initial loss is set to infinity); [2] 360 epochs have been finished. Training was done using keras (Chollet and others, 2015) with the theano backend (The Theano Development Team *et al.*, 2016).  The final MAPE and MSE in Figure 4 were tested on the testing dataset. |
| Derivation of model parameters for PWMs transformed by the exact transformation | This is due to Theorems 1 and 2. Below is an effort to make it clear for practice.  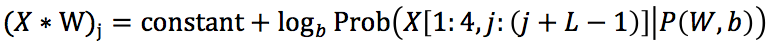We noted that Theorem 1 has the following form:  For the original CNN model, we have the following linear transformation l with the matrix of max-pooling of convolutional values **C** (n samples (rows) and K kernels/PWMs (columns)):  $l\left( \mathbf{C} \right)\mathbf{= C}\cdot\mathbf{H}+\boldsymbol{1}_{n}\cdot\mathbf{z}^{T}$  where **H** (k rows and p columns) and **z** (vector of length p) are weights and biases for the linear transformation.  By Theorem 1 we have  $\mathbf{C}=\boldsymbol{1}_{n}\cdot\mathbf{d}^{T}+(\log_{b} b')\cdot\mathbf{L}$  where **L** is the matrix of maximal log-likelihood values (n rows and K columns; with an arbitrary base of logarithm b’) and **d** the vector of constants for all kernels (thus with length K).  Inserting this into the linear transformation above gives us:  $l\left( \mathbf{L} \right)\mathbf{=}\left( \boldsymbol{1}_{n}\cdot\mathbf{d}^{T}+(\log_{b} b')\cdot\mathbf{L} \right)\cdot\mathbf{H}+\boldsymbol{1}_{n}\cdot\mathbf{z}^{T}\boldsymbol{=}(\log_{b} b^{'})\cdot\mathbf{H}\cdot\mathbf{L+}\boldsymbol{1}_{n}\cdot{(\mathbf{H}^{T}\boldsymbol{\cdot d+z})}^{T}$  Therefore, if we apply a linear regression on **L**, with weights $(\log_{b} b^{'})\cdot\mathbf{H}$ and biases $\mathbf{H}^{T}\boldsymbol{\cdot d+z}$ , we will get the same result as $l\left( \mathbf{C} \right)\mathbf{= C}\cdot\mathbf{H}+\boldsymbol{1}_{n}\cdot\mathbf{z}^{T}$. This is how we derive the appropriate model parameters. |
