## Supplementary Table 3 for "An exact transformation for CNN kernel enables accurate sequence motif identification and leads to a potentially full probabilistic interpretation of CNN"

Supplementary Table 3. Details for the AddGene analysis

| Dataset description | A JSON file describing the sequence and lab-of-origin of each plasmid.  Requested from AddGene ([www.addgene.org](http://www.addgene.org)).  The original version of this dataset used by the original authors (Nielsen and Voigt, 2018) is not directly available from the authors or AddGene itself; instead, we received an augmented version of this dataset from AddGene. This version has 72594 samples (before preprocessed by the following scripts) / 64149 samples (after preprocessed by the following scripts) and is available upon request. |
| --- | --- |
| Preparation of CNN model, kernel-to-PWM transformation, and final evaluation | Scripts are downloaded from the Git repository: <https://github.com/VoigtLab/predict-lab-origin> . We used the commit a4c641c and adapted some changes to make the scripts free of error.  Because the AddGene dataset has been augmented (as described above), we need to retrain the CNN model first before evaluating the transformations.  Details about code adaptation, CNN retraining, kernel-to-PWM transformation and final evaluation (and the evaluation results) are stated in the following file. Note that, because there are N’s in input sequences, we need to compute extra constants to make the exact transformation work (see Section “Calculate the (corrected) log-likelihoods” in the README.md below).  https://github.com/gao-lab/kernel-to-PWM/tree/master/Supplementary_code_for_manuscript/ work06_compare.k2p.and.Deepbind.heuristic.for.complex.models/README.md |
